# Hox-A2 protein expression in avian jaws cartilages and muscle primordia development

**DOI:** 10.1101/2024.03.14.584976

**Authors:** Stéphane Louryan, Myriam Duterre, Nathalie Vanmuylder

**Affiliations:** Laboratory of Anatomy, Biomechanics and Organogenesis, Faculty of Medicine, Université Libre de Bruxelles, route de Lennik, 808 (CP 619), B1070 Bruxelles (Belgique)

**Keywords:** Hox-A2, chick embryo, cephalic cartilages, muscles

## Abstract

Objective: to elucidate the branchial origin of the articular and the square (homology of the malleus and the incus of mammals), we used immunohistochemistry to analyse the expression of the Hox-A2 protein during cephalogenesis in chickens.

Materials and methods: immunohistochemistry on paraffin sections of embryos from stage HH16 to HH40.

Results: in addition to the columella (equivalent to the mammalian stapes), the joint between the articular and the quadrate bones, and the retroarticular process of the articular (homologous to the short process of the malleus) express Hox-A2, suggesting an intervention of the 2^nd^ arch in their formation. However, we fortuitously observed very intense expression within the early muscle plate of the second arch, which then generalized to all cephalic muscles, and extended to the trunk’s myotomes. In the cartilage, the presence of the protein disappeared at stage 35.

Discussion and conclusion: the present results, while confirming the contribution of the second arch to the development of avian equivalents of the mammalian ear ossicles, strongly suggest that the Hox-A2 gene plays a role in muscle development, which remains to be elucidated by more sophisticated techniques.

## Introduction

The branchial origin of mammalian middle ear ossicles has been known for numerous years. However, the precise contribution of the first (mandibular) and second (hyoid) arch is still discussed [1]

The classical and more accepted theory implies that malleus and incus arise from the first pharyngeal arch cartilage (Meckel’s cartilage), and the stapes from the second one (Reichert’s cartilage). This hypothesis is validated by some morphologic evidence [2] and is consistent with several experimental data relative to Hox genes knock-out and overexpression [3, 4]

However, another model could be proposed, based upon morphologic features and in vitro experiments, in which the ventral parts of the malleus (including the handle) and incus derive from the second arch cartilage, with the stapes and the laterohyale [5–14]

To decide between these two hypotheses, Louryan et al. [1] studied the expression of Hox-A2 protein using immunohistochemistry during mouse ossicle development. HOXA2 gene is known to express in second-arch derivatives, but not in the first-arch elements.

Surprisingly, every ossicle anlage contained Hox-A2-positive cells, in variate proportions, even if the main Hoxa-2 expression was observed in the Reichert’s cartilage, the stapes rudiment and the short process of the malleus. This observation suggested that second arch-derived cells were present in the entire ossicular chain, even if the main derivatives of the second arch are the stapes and the short process of the malleus.

In terms of phylogenetic evolution, the Reichert-Gaupp theory postulates that the malleus is the remnant of the articular bone of reptiles and birds and that the incus derives from the quadrate bone. The stapes corresponds to the columella. The handle of the malleus was initially associated with the retro-articular process of the articular bone [15–17]. However, more recent data trend to associate more precisely the short process of the malleus with the retro-articular process. Indeed, the more ancient investigations did not distinguish between the handle (considered as the part of the malleus connecting with the eardrum) and the short process, which suggests taking into account questions of terminology and the fact that the anatomy of the malleus may differ among species.

Then, if the developmental features of homologous elements are similar, it could be possible to find Hoxa-2-positive cells in several components of articular and quadrate bones.

The initial purpose of this study was to analyse Hox-A2 protein expression in the rudiments of quadrate and articular bones of the chicken embryo, with consideration for the surrounding tissues. However, some unusual observations during our investigation led us to make further observations, particularly relating to muscular development.

### Material and methods

Embryonic stages were established following Hamilton [18]. The more adequate stage for cartilage development was determined following Vorster [19]. Stages 16, 17, 21, 24, 27 embryos were removed from the egg. They were fixed in Serra’s fixative medium (Ethanol 95%: 6 parts; formalin 40%: 3 parts; acetic acid 100%: 1 part), dehydrated in a graded series of alcohol, and embedded in paraffin. 5 µm transverse or sagittal sections were placed on slides for staining and immunohistochemistry. Alternate slides were routinely rehydrated and stained with Hematoxylin-Eosin or Masson’s trichrome. For antigen retrieval, the slides were placed in citrate buffer in a microwave for 20 minutes. Sections were washed in distilled water for 5 minutes, in PBS 0.01M containing 0.1% Triton to permeabilize the cytoplasmic membrane for 5 minutes, then three times 5 minutes in PBS. They were submitted for 1 hour to blocking serum (Product 20773, EMD Millipore Corp., Burlington, MA, USA).. They were then incubated overnight in a humidified chamber with 1/100 rabbit anti-HOXA-2 (orb 184090, Biorbyt, Cambridge, UK). Endogenous peroxidases were blocked in methanol H_2_O_2_ 0,3 % for 30 minutes. The secondary biotinylated goat anti-mouse IgG (BA-9200, Vector Laboratories, Burlingame, California, USA) antibody was diluted 1/200, added to the samples, and incubated for 30 min. They were incubated for 10 min in the presence of Streptavidin (Vectastain Elite ABC kit Peroxidase (HRP, Vector Laboratories, Newark, Ca, USA), then stained by DAB (HIGHDEF DAB, product ENZ-ACC105-0200, Enzo Life Science, NY, USA).

Negative controls were performed by omitting the first antibody.

Ethical statements: following the European and Belgian laws, “non-free-living larval forms” sampling does not require ethical committee advice.

## Results

At HH 16 stage (Fig. 1), there was not yet precartilaginous primordia. The visceral arches were quite present. Surprisingly, a very strong immunoreactivity appeared in the premuscular plate of the second arch in comparison to the first arch one, which was less intensively labeled (Fig. 1A and B)

**Fig. 1:**
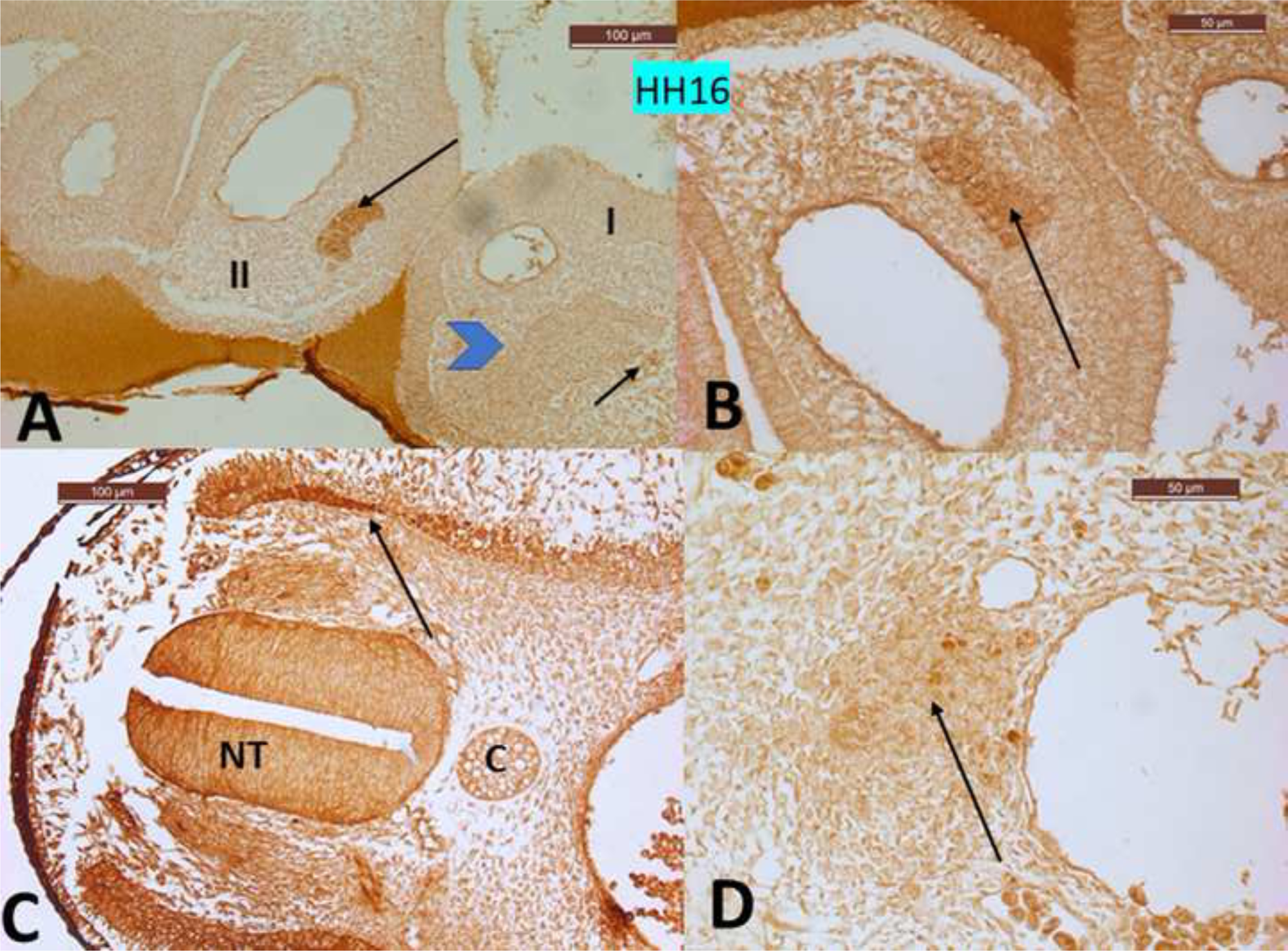
HH 16 embryo. A and B: intense immunostaining of Hox-A2 in the hyoid muscle plate (arrows). In A, the mandibular muscle plate (large arrow) appears globally not labeled, but several peripheral cells appear positive (arrow at the right side). I: first arch, II: second arch. C: numerous positive cells in the trunk myotomes (arrows). NT: neural tube. D: some cells of oculomotor muscle plate are labeled (arrow). All sections are in the sagittal plane except C (transverse section).

A similar immunoreactivity was observed in in the trunk myotomes (Fig. 1C) as well in the premuscular oculomotor primordium (Fig. 1D).

At the HH 17 stage (Fig. 2), the muscular plates were dissociated, and first and second-arch immunoreactive cells appeared sparse in the mesenchyme The labeled cells were more numerous in the second arch (Fig. 2B). This observation was the same in the trunk myotomes. In the oculomotor area, we observed persistent positive cells.

**Fig. 2:**
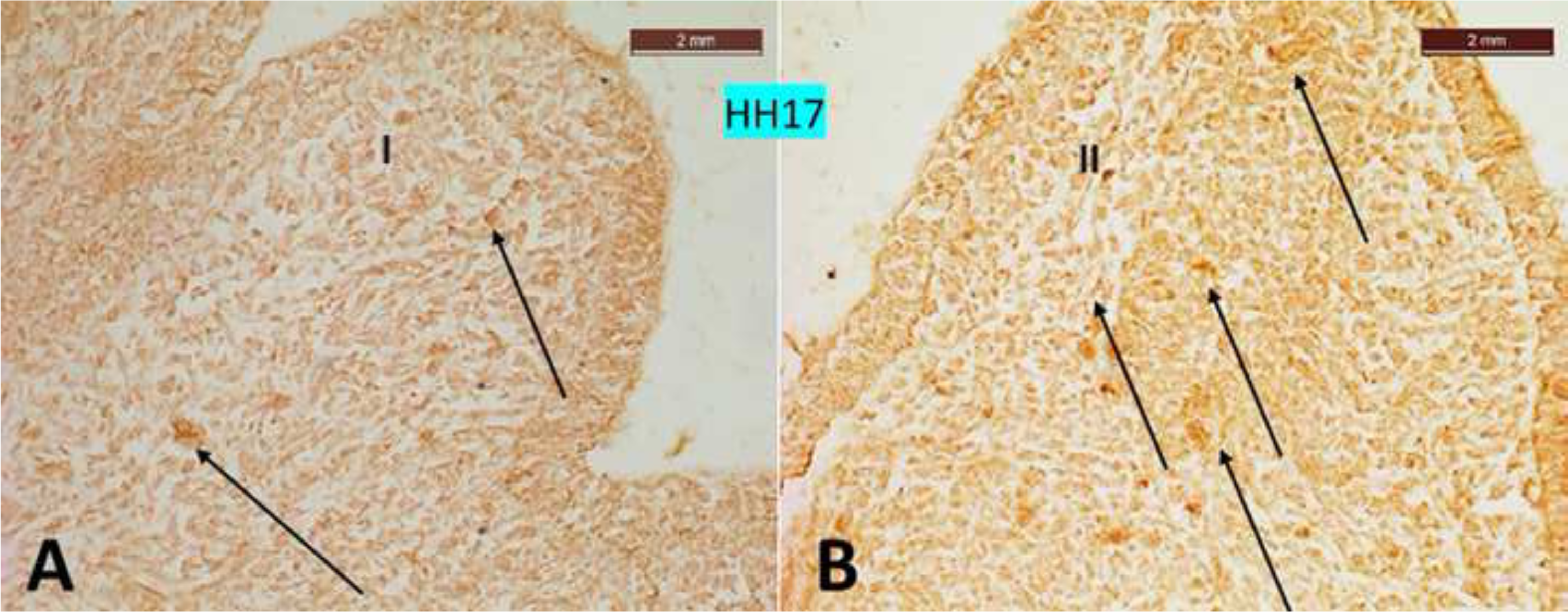
HH17 embryo, sagittal sections. Comparison between Hox-A2 staining in the premuscular cells in the first (A; I) and second (B; II) arches. The cells are more numerous in the second arch, in which the muscle plate is still recognizable.

At HH21 and 24 stages (Fig. 3 and 4), the observations were similar. A population of intensely stained cells remained visible in the future area corresponding to oculomotor muscles (Fig. 4A). In the trunk somites, the migration pathways of premuscular cells were clearly distinguished (Fig. 4B).

**Fig. 3:**
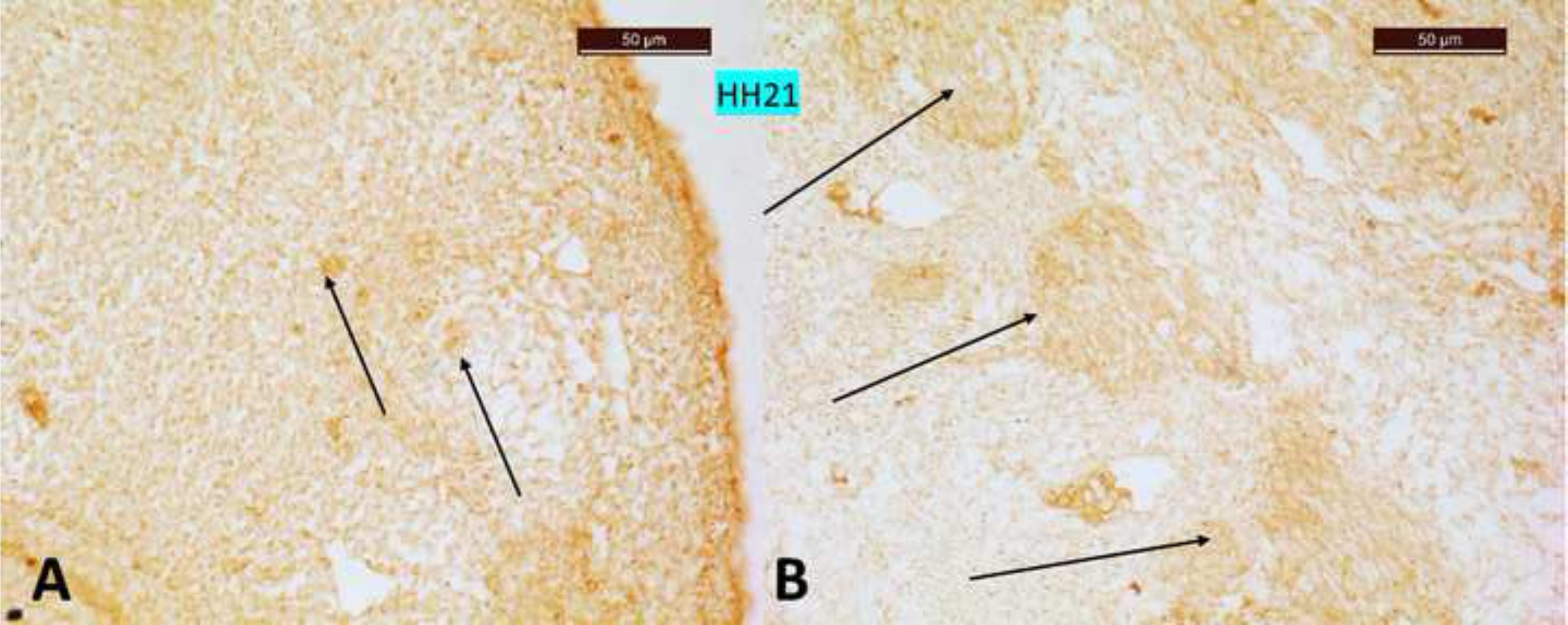
HH 21 embryo, sagittal sections. A: persistence of sparsely labeled cells (arrows) in the second arch. B: segmented groups of labeled cells in the trunk myotomes (arrows).

**Fig. 4:**
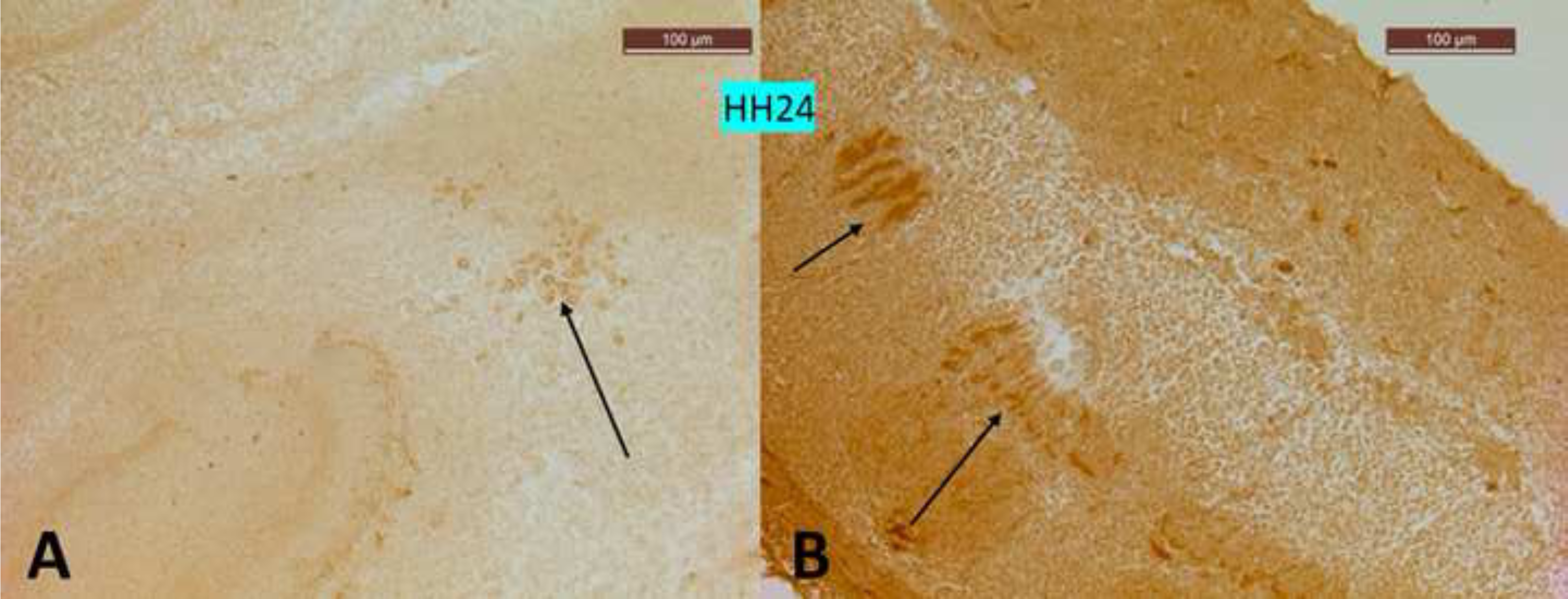
HH24 embryo, sagittal sections. A: group of positive cells in the oculomotor muscle plate (arrow); B: intense staining in the metameric trunk myotomes (arrows).

From the HH 27 stage (Fig. 5), the precartilaginous blastemata became apparent. Immunoreactive cells were essentially present in the perichondrium but concerned also chondroblasts. Positive cells predominated in the columella auris, in the interzone between articular and quadrate bones, and the retro-articular process of the articular (Fig. 5A to C). The myotubes of the depressor and adductor mandibulae muscles were also strongly stained (Fig. 5B and C). Reichert’s cartilage was intensely stained contrary to Meckel’s one, but the surrounding muscular primordia were equally stained around both of them (Fig.5D and E). A diffuse immunoreactivity is visible in the mesenchyme around the Reichert’s cartilage and the second branchial arch muscles precursor, and is absent in the first arch area. The oculomotor muscles began to differentiate and were stained (Fig.5F). Trunk myogenic cells remained positive too.

**Fig. 5:**
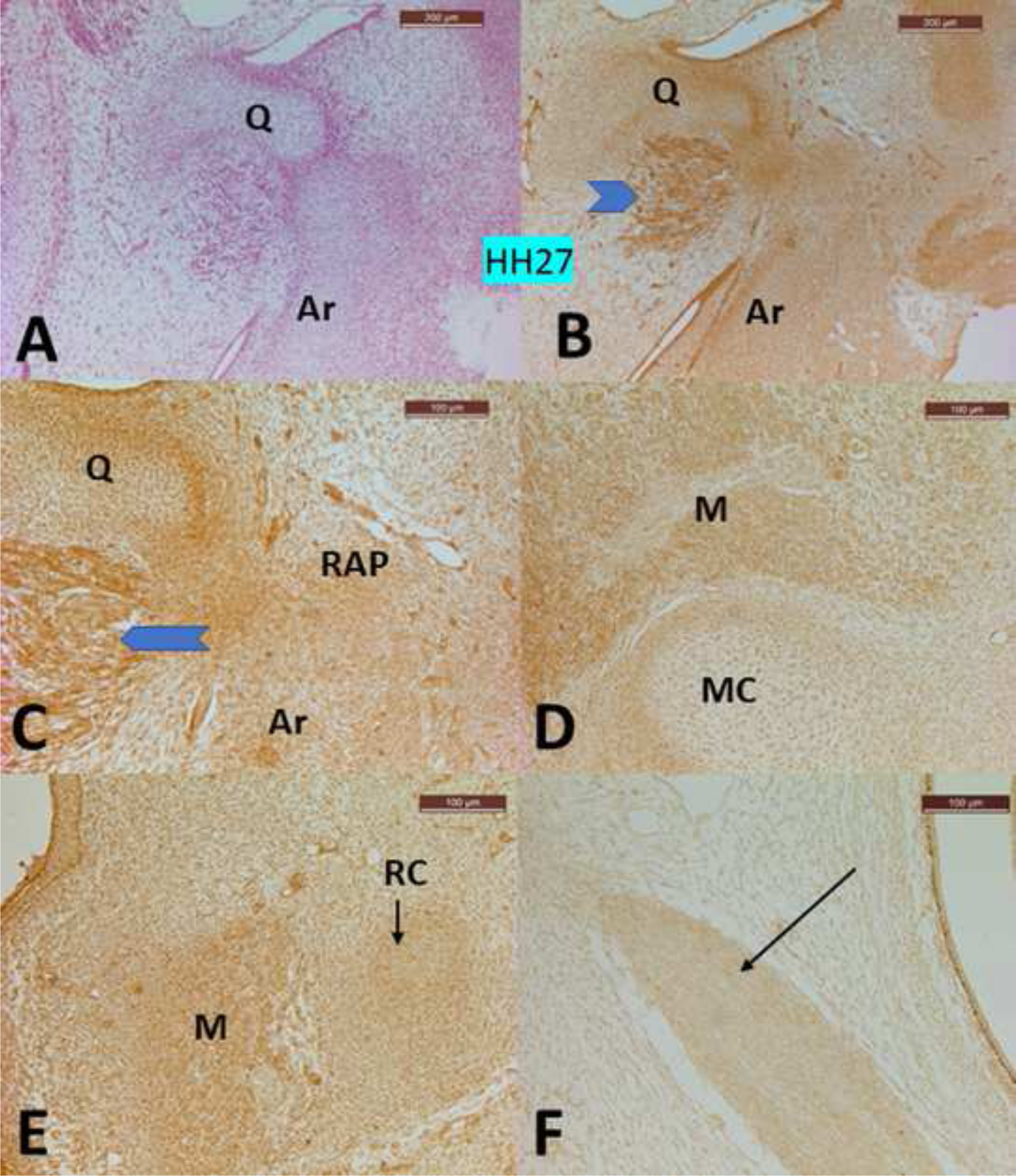
HH27 embryo, sagittal sections. A (Hematoxylin-Eosin), B, and C: articular (Ar)/quadrate (Q) cartilages articulation. The Hox-A2 staining is present in the periphery of both cartilages in the articular zone, but also in the retro-articular process (RAP). An intense labeling appears in the depressor mandibulae muscle primordium (large arrow). D: section in the first arch: the Meckel’s cartilage is unstained but surrounding muscles (M) are labeled. In the second arch (E), the Reichert’s cartilage appears positive as the muscle (M) mass. F: positive rectus inferior muscle.

At HH 29 stage (Fig. 6), we observed the persistence of the same staining, but there was a strong difference between the articular-quadrate complex, in which the staining is essentially peripheral, and the strong and diffuse immunoreactivity observed in the columella and the Reichert’s cartilage.

**Fig. 6:**
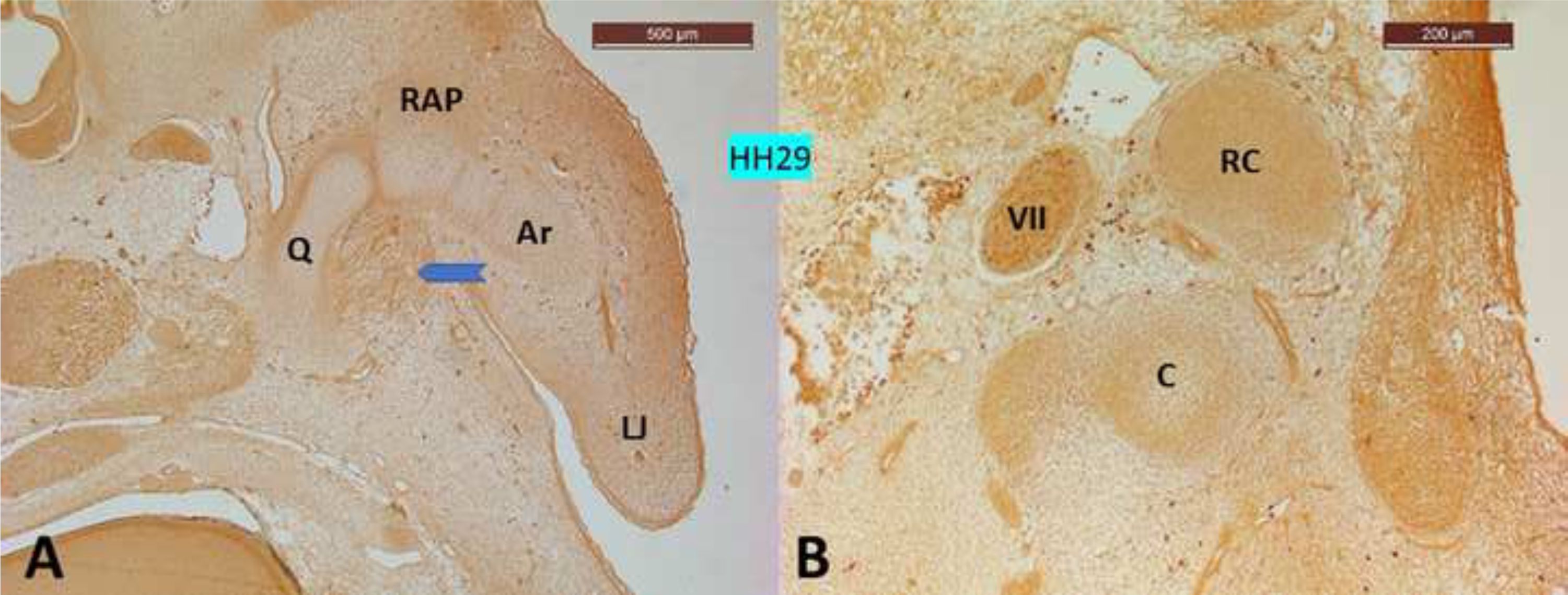
HH27 embryo, sagittal sections. A: articular (Ar)/quadrate (Q) cartilages articulation; similar findings than the previous figure. LJ: lower jaw. Other captions: see fig. 5. B: peripheral labeling of the columella auris © and massive staining in the Reichert’s cartilage. The facial nerve (VII) is also labeled.

At HH35 and 40 stages (Fig. 7 and 8), the cartilage immunoreactivity has disappeared, in the columella as well in the retro-articular process. However, all muscle primordia were positive, Including the entire oculomotor primordia.

**Fig. 7:**
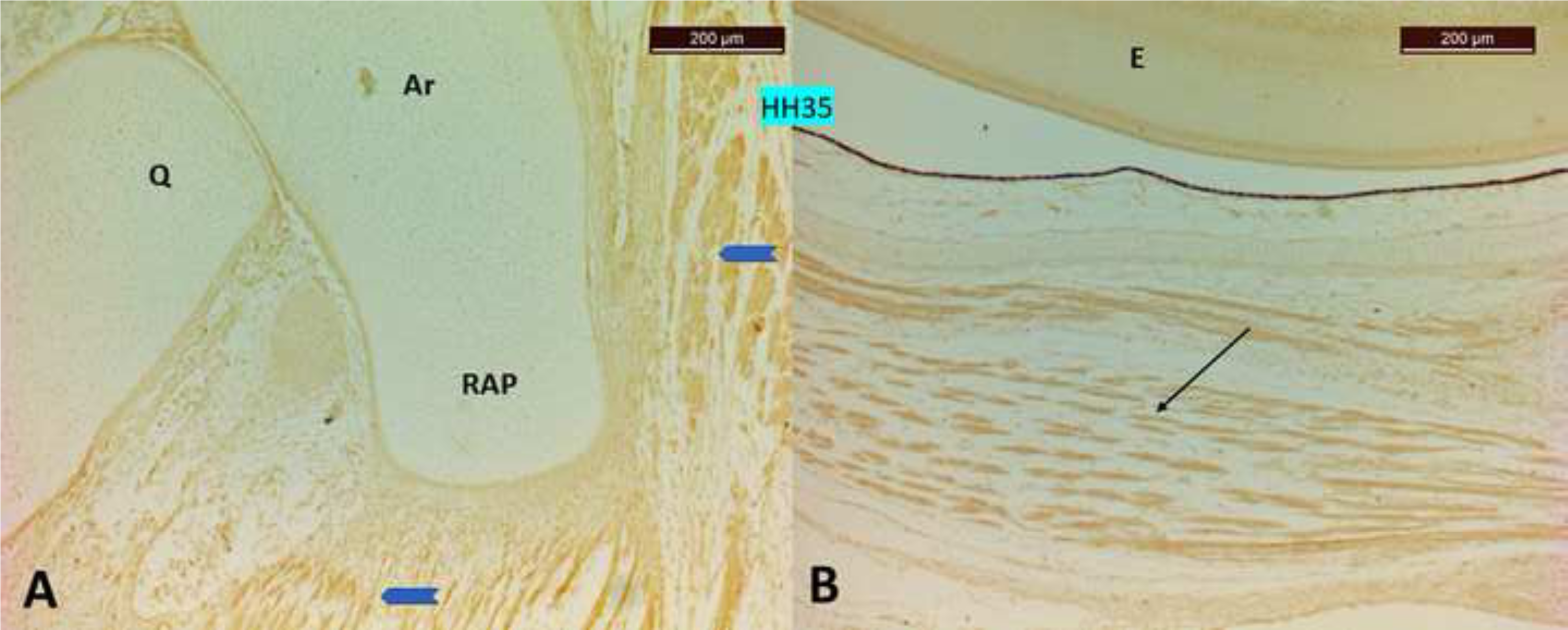
HH35 embryo, sagittal sections. A: articular (Ar)/quadrate (Q) cartilages articulation. At stage HH35, the cartilages are no longer labeled, at contrary to the muscle primordia (large arrow). For captions, see Fig. 5. B: labeled rectus inferior muscle.

**Fig. 8:**
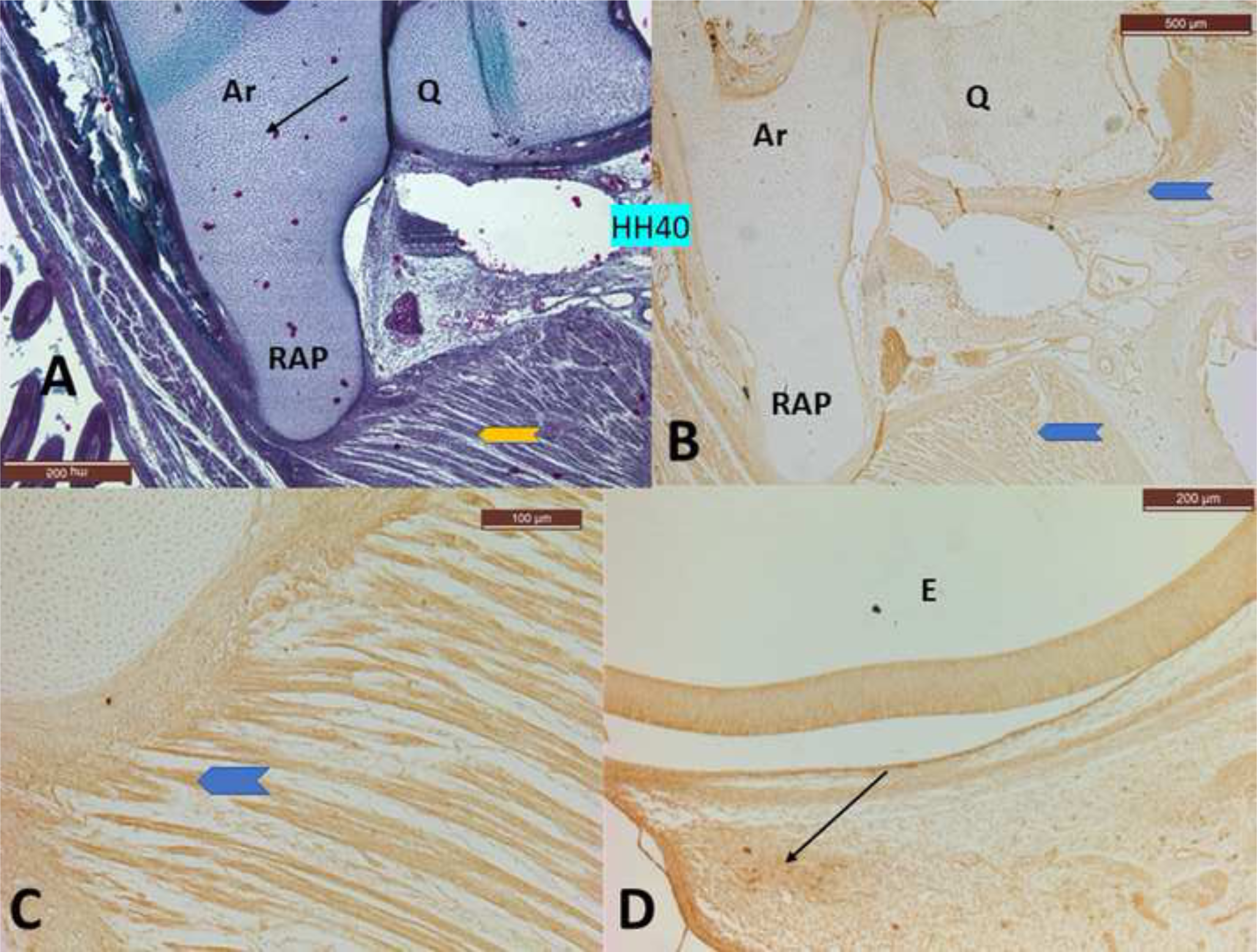
HH40 embryo, sagittal sections: A, B, C: similar features as Fig. 7 (A: toluidine blue). The muscular primordia are labeled using large arrows. Same captions as the previous illustrations. D: intense staining in the depressor of the lower palpebra primordium (arrow)

Negative control did not exhibit any staining.

## Discussion

HOXA2 is known to play an important role in patterning the second pharyngeal arch in the embryo; it constitutes a key factor for the second arch derivatives, as the Reichert’s cartilage, the stapes (columella in the bird) and associated structures. It is not expressed in the first arch [20–23].

Its expression in cartilages predominates in the peripheral parts of them [20]. However, it has been demonstrated that Hox-A2 is expressed in the mouse palatal primordia, independently from the second pharyngeal arch, and that this gene could play a role in the muscular connections in the Xenopus [24] Concerning the mammalian middle ear ossicles, Louryan et al. [1] showed that, if Hox-A2 expressed strongly in the stapes, Reichert’s cartilage, and the short process of the malleus, a diffuse immunoreactivity was observed in the whole ossicular primordia, while the Meckel’s cartilage of the first arch displayed no staining. This confirms the close association of stapes end short process of the malleus with the second arch but suggests a wider contribution of the second arch to the ossicular chain, already supported by morphologic and experimental previous studies [5–14]. Using in situ hybridization, Santagati et al. [22] published interesting pictures in which HOXA was present in the malleus and the incus, without comment on this unexpected observation. The fact that a transient immunoreactivity was observed in articular and quadrate bone, with a maximum intensity in the retro-articular process of the articular reinforces previous suggestions about the contributions of cells arising from the second arch and confirms the homology between the retro-articular process and the short process of the incus (which takes part of the handle of the malleus). However, in the early stages, immunohistochemistry does not identify massive staining in the undifferentiated second arch mesenchyme as shown using in situ hybridization [20].

Because palatal primordia express Hox-A2 [23], without any relationship with the second arch, it should be possible that Hox-A2 staining occurs on some cell populations independently of their branchial origin, and that its expression could not be considered a reliable marker to solve the problem relative to the contribution of each arch to the bone development.

If we now consider the cephalic muscles, the fact that Hox-A2 is very strongly expressed in the hyoid muscle plate is consistent with our knowledge about the close relationship of this gene with the second arch. The first arch muscle plate was poorly stained and later, the staining extended to all premuscular and muscular primordia whatever their origin and location, including trunk myotomes and muscle rudiments. This suggests a key role of HOXA2 in muscle differentiation, regardless of their locations.

In the earlier stages of their development, however, only second arch-derived cephalic pre-myoblast strongly express the gene, including some cells destinated to the oculomotor muscles, part of which derives from the amniote and teleost 5^th^ somitomere, which develops in front to the second and third rhombomeres, in which lies the more cranial expression of HOXA2 in the neural tube.

The 4^th^ and 5^th^ amniote and teleost somitomeres correspond to the selachian and amphibian third one. This 5^th^ somitomere is known in the chick embryo to give rise to the lateral rectus and palpebral depressor muscles [25,26]

In the selachians embryos, there seems to exist a morphological continuity between the cavities of the branchial muscle plates (myocoeles) and the corresponding oculomotor primordia, around the dorsoventral axis [27–29], each of them constituting the dorsal (for instance innervated by nerve VI) and ventral parts (depending on nerve VII) of common somitic anlagen. This question is largely discussed by Kuratani and Adachi [30]. So, Brachet [31] and De Beer [32] associated the blastema of the rectus externis muscle with the “hyoid somite” (corresponding in fact to the third urodelian and selacian somitomere)-also responsible for the muscles of the hyoid arch.

The evolution of myogenic cells arising from the muscle plate, with progressive dissociation, is similar to the findings of Milaire (1999) using ATP-phosphohydrolase staining in the chick embryo [33].

In summary, the premuscular primordia directly or indirectly associated with the hyoid arch and the trunk myotomes express early and strong HOXA2. Later, more differentiated muscular elements become immunoreactive, without considering their origin. This suggests a key role of Hoxa-2 in muscle patterning and constitutes a new function to add to the multiple properties of this gene. It is possible that precocious strong expression of the gene in the hyoid muscle plate, followed by larger staining in muscular primordia, could correlate with the precocious deep relationship between this gene and the second pharyngeal arch, with as a consequence an earlier premuscular differentiation. Further experiments using in situ hybridization should be performed to refine these preliminary results, particularly to elucidate the pathways involved in the Hox-A2 function on muscular development.

